# Vast differences in strain-level diversity in the gut microbiota of two closely related honey bee species

**DOI:** 10.1101/2020.01.23.916296

**Authors:** Kirsten M Ellegaard, Shota Suenami, Ryo Miyazaki, Philipp Engel

## Abstract

Most bacterial species encompass strains with vastly different gene content. Strain diversity in microbial communities is therefore considered to be of functional importance. Yet, little is known about the extent to which related microbial communities differ in diversity at this level and which underlying mechanisms may constrain and maintain strain-level diversity. Here, we used shotgun metagenomics to characterize and compare the gut microbiota of two honey bee species, *Apis mellifera* and *Apis cerana,* which have diverged about 6 mio years ago. While both host species are colonized by largely overlapping bacterial 16S rRNA phylotypes, we find that their communities are highly host-specific when analyzed with genomic resolution. Despite their similar ecology, *A. mellifera* displayed a much higher extent of strain-level diversity and functional gene content in the microbiota than *A. cerana,* per colony and per individual bee. In particular, the gene repertoire for polysaccharide degradation was massively expanded in the microbiota of *A. mellifera* relative to *A. cerana*. Bee management practices, divergent ecological adaptation, or habitat size may have contributed to the observed differences in microbiota composition of these two key pollinator species. Our results illustrate that the gut microbiota of closely related animal hosts can differ vastly in genomic diversity despite sharing similar levels of diversity at the 16S rRNA gene. This is likely to have consequences for gut microbiota functioning and host-symbiont interactions, highlighting the need for metagenomic studies to understand the ecology and evolution of microbial communities.

## Introduction

Most bacteria live in complex communities, composed of hundreds to thousands of species, which in turn encompass strains with highly variable gene content [1, 2]. Also in host-associated bacterial communities, strain-level diversity can be substantial, despite the general assumption that genetic diversity destabilizes mutualistic interactions [3]. For example, multiple strains of a sulfur-oxidizing endosymbiont were found to co-colonize individual hosts of deep-sea mussels, presumably because they encode complementary functions [4, 5]. In contrast, strains of the human gut microbiota have been shown to segregate among individuals, resulting in host-specific genetic profiles [6–8]. However, despite the increased awareness of the existence and functional importance of strain-level diversity in host-associated bacterial communities, little is known about differences in diversity across host species, or the underlying mechanisms which constrain and maintain diversity within and among hosts. This is largely due to (i) the limited insights gained from 16S rRNA gene amplicon sequence datasets about strain-level diversity [9–11], (ii) the technical challenges associated with the deep-sequencing and analysis of bacterial genomes from complex natural communities [12], and (iii) the difficulty to generate comparable datasets across host organisms.

Unlike most animals, eusocial corbiculate bees (honey bees, stingless bees, and bumble bees) have been shown to harbor a specialized gut microbiota with a simple and highly consistent taxonomic composition, consisting of up to 10 phylotypes, as based on 16S rRNA gene analyses [13]. Considering the common occurrence of these phylotypes across bees, it is plausible that they were acquired around the time when eusociality evolved in the bees [13]. Interestingly, previous studies on bacterial isolates have demonstrated an impressive amount of genomic diversity within phylotypes, where strains isolated from different hosts often represent divergent phylogenetic sub-lineages [14–17]. Moreover, a recent metagenomic analysis of the gut microbiota of the Western honey bee, *Apis mellifera,* found that even within the same host species divergent sub-lineages can be present [6]. We refer to these sub-lineages as sequence-discrete populations (SDPs), since metagenomic data demonstrated that they were discrete from each other [6], in addition to being sufficiently divergent to be considered as different bacterial species [18]. Given these results, it is possible that bee species with similar phylotype-level composition in their gut microbiota harbor very different communities. Indeed, two previous studies based on amplicon sequence data have provided first evidence that the extent of diversity can differ between different species of social bees [13, 19]. However, comparative community-wide analyses, based on data with genome-level resolution, are currently lacking.

In the current study, we perform a comparative metagenomic analysis of the gut microbiota of two closely related species of honey bees, *Apis mellifera* and *Apis cerana*. Based on molecular data, their last common ancestor has been dated to approximately 6 million years ago [20–23], and previous 16S rRNA-based studies have shown that they harbor largely overlapping gut microbiota phylotypes (>97% 16S rRNA identity) [13]. With the exception of *A. mellifera*, all extant species of honey bees (genus *Apis*) are confined to Asia, pointing towards an Asian origin of the *Apis* genus [21]. Based on molecular analysis, *A. mellifera* expanded into its native range (Africa, Europe, and Western Asia) approximately 300.000 years ago [20]. However, *A. mellifera* was recently re-introduced to Asia by humans [24, 25], thereby bringing not only the bees, but also their associated bacterial communities into close proximity, potentially resulting in a homogenization of their gut microbiota.

In order to compare the composition, diversity, and evolution of the gut microbiota of *A. mellifera* and *A. cerana* at the strain-level, we analyzed shotgun metagenomes of individual bees, using a common DNA extraction protocol and comparable sequencing depth. We find that each host species harbors a highly distinct bacterial community, largely composed of different SDPs and strains, with occasional transfers among sympatric bees. Quantitative analysis revealed that the gut microbiota diversity of *A. mellifera* is much higher than for *A. cerana*, resulting in a larger metabolic flexibility at both the individual and colony level. These results represent the first comparative genome-wide analysis of strain-level diversity between related host-associated microbial communities, raising new questions regarding underlying mechanisms and functional consequences.

## Results

### Metagenomic data reveals that the gut microbiota of *A. mellifera* and *A. cerana* are distinct

A total of 40 shotgun metagenome samples were collected from individual bees, with 20 bees per host, all representing nurse bees that were sampled from inside the hive. Two colonies were sampled from each host, all from different apiaries, no more than 100km apart, close to Tsukuba (Japan). In order to perform a metagenomic characterization of the community composition across samples, we first established a genomic database of isolated strains (**Dataset S1**) representative of both hosts. In total, 10 new genomes isolated from *A. cerana* were sequenced and added to a previously established, non-redundant honey bee gut microbiota database [6], together with previously published genomes isolated from more distantly related bee species. Approximately 90% of the host-filtered reads mapped to the new database, regardless of the host affiliation of the samples, indicating that the database is highly and equally representative of the gut microbiota of both hosts (**Fig. S1**).

To make an initial broad comparison of the gut microbiota composition, the relative abundance of all phylotypes in the database was quantified, based on mapped read coverage to single-copy core genes. Overall, the phylotype-level composition was largely consistent with previous studies employing amplicon sequencing of the 16S rRNA gene [13]. The five phylotypes which have been proposed to constitute the core microbiota of corbiculate bees were found to colonize both hosts, although the relative abundance profiles were distinct between the hosts (**Fig. S2**). Other phylotypes associated with *A. mellifera* (*Bartonella apis*, *Frischella perrara*, *Commensalibacter* sp.) were not detected in any of the *A. cerana* samples, consistent with a very low prevalence in this host[13] Conversely, while *Apibacter* was not detected in any of the *A. mellifera* samples, it was prevalent among the *A. cerana* samples (**Fig. S2**, yellow color).

To determine whether the five core phylotypes colonizing both hosts are distinct at the SDP level, candidate SDPs were inferred from isolate genomes in the database, based on core genome phylogenies (**Fig. 1A-C,G,H**) and pairwise average nucleotide identities (ANI) (**Dataset S2**). Subsequent metagenomic validation (**Fig. S3-S4**) confirmed three new SDPs within the *Gilliamella* phylotype (**Fig.1A**), and one new SDP within the *Lactobacillus* Firm5 phylotype (**Fig. 1B**), all of which were represented exclusively by *A. cerana*-derived isolates. A new SDP was also confirmed for *Snodgrassella* (**Fig. 1C**), based on isolates from *A. cerana*, *A. andreiformis* and *A. florea*, suggesting that this SDP may be shared among other species of honey bees than *A. mellifera*. In contrast, for the two remaining core phylotypes *Lactobacillus* Firm4 and *Bifidobacterium*, no new candidate SDPs were inferred (**Fig. 1G-H**), since the *A. cerana*-derived genome isolates had ANI values of up to 95% and 90% to the *A. mellifera* isolates, thus falling within the range of ANI values observed among the *A. mellifera* isolates (**Dataset S2**).

**Figure 1.**
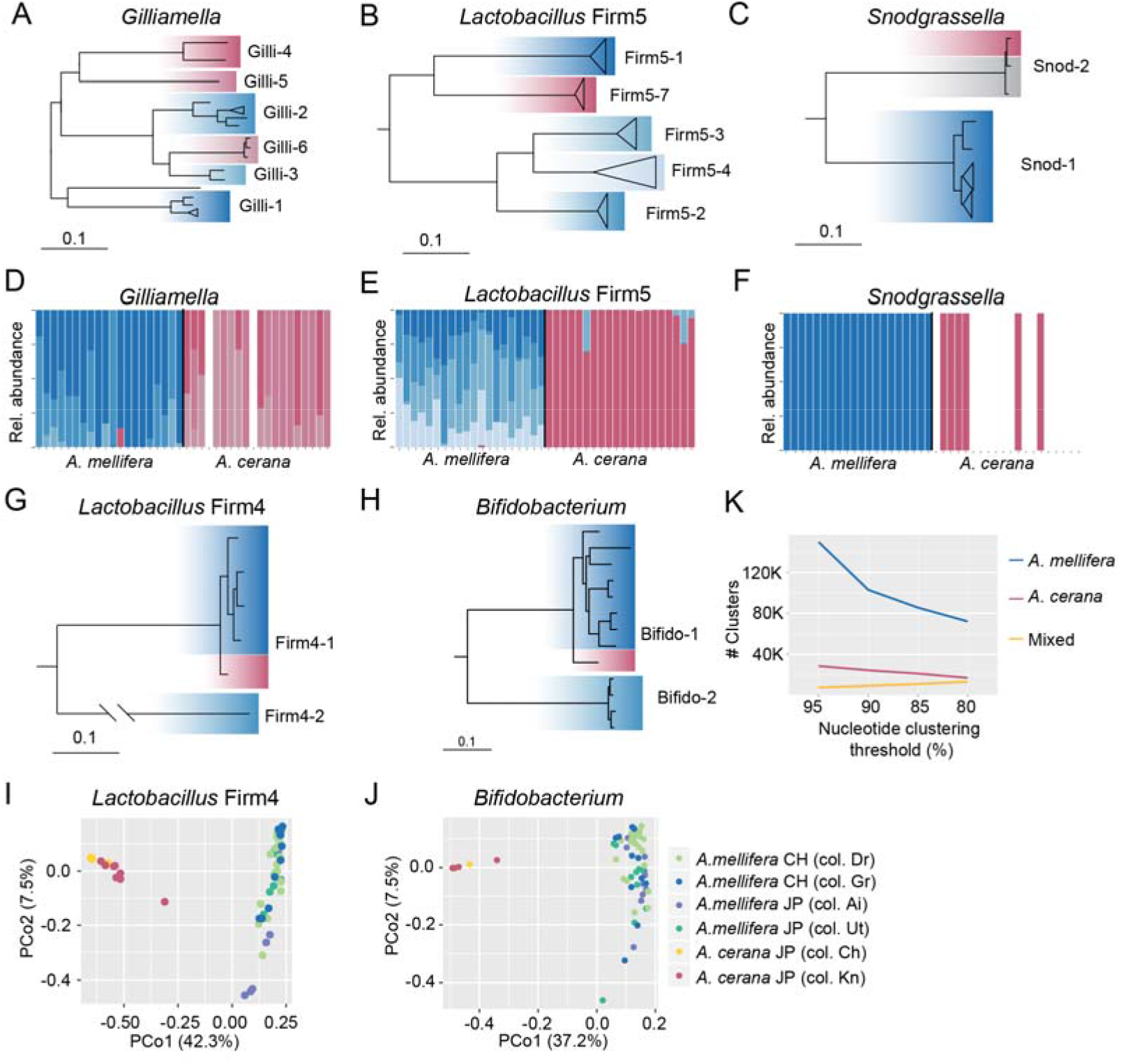
The gut microbiota of *A. mellifera* and *A. cerana* are composed of divergent SDPs and strains. **(A-C, G-H)** Core genome phylogenies of the five shared core phylotypes colonizing *A. mellifera* and *A. cerana*. Confirmed SDPs are indicated by the labels of the clades. Genomes are highlighted with blue shades for isolates from *A. mellifera*, and red shades for isolates from *A. cerana*. Grey shades indicate isolates from other honey bee species. Bars correspond to 0.1 substitutions per site **(D-F)** Barplots displaying relative abundance of the confirmed SDPs shown in panels A-C, across metagenomes from Japan (see Fig. S5 for the same plots including metagenome samples from Switzerland). **(I-J)** PCoA ordination plots based on the pairwise fractions of shared SNVs (jaccard distance) (see **Dataset S3** for plots of other SDPs). Dots represent individual samples, color-coded by host and colony origin as indicated by the legend. **(K)** Number of host-specific and mixed sequence clusters generated from metagenomic ORFs at different clustering thresholds for the complete dataset (Swiss and Japanese samples).

To further validate the host specificity of the novel SDPs and detect eventual cross-host transfers, we quantified the relative abundance of each SDP across the metagenomic samples, including samples previously collected in Switzerland. All SDPs found to be host-specific in the genomic database displayed a clear host preference across the metagenomic samples, but a small number of transfers were nevertheless detected among the Japanese samples within the *Lactobacillus* Firm5 and *Gilliamella* phylotypes (**Fig. 1D-F**). These results therefore indicate that honey bees do get exposed to non-native SDPs, at least in Japan, occasionally resulting in colonization. For the SDPs previously described for *A. mellifera* [6], the abundance patterns were very similar between Swiss and Japanese samples, with co-occurrence of SDPs within individuals being the norm for all core phylotypes except *Gilliamella* (**Fig. 1D-F** and **Fig. S5**). Thus, these distributions likely represent conserved patterns of co-existence.

For the two core phylotypes harboring SDPs with genome isolates from both hosts (**Fig. 1G-H**), rooting of the core genome phylogenies with related SDPs showed that the *A. cerana*-derived isolates diverged prior to the *A. mellifera*-derived isolates, as would be expected if a more recent host-specialization had occurred. Therefore, to determine whether the communities differ at the strain-level, the fraction of shared single-nucleotide variants was calculated for all sample pairs, and visualized using principal coordinate analysis (**Fig. 1I and J**). Remarkably, the samples were found to cluster strongly by host, indicating that each host is colonized by a distinct population of strains. In contrast, there was no clustering by country or colony affiliation (**Fig. 1I-J**). For other community members, clustering by country or colony was observed for only a subset of SDPs (**Dataset S3),** indicating that strains are not necessarily geographically specialized.

Finally, to obtain a database-independent estimate of the divergence between the gut microbiota of *A. mellifera* and *A. cerana*, we *de novo* assembled the metagenomes and compared the gene contents by clustering all predicted ORFs (open reading-frames) by sequence identity. Metagenomic assemblies and ORFs were inferred individually for each sample, using a subset of 20 million paired-end host-filtered reads per sample, and clustering was done using a range of thresholds (80-95% nucleotide identity). Only a very minor fraction of the clusters contained sequences from both hosts (**Fig. 1K**) further corroborating the small overlap of the gut microbiota between the two honey bee species.

In conclusion, although the gut microbiota of *A. mellifera* and *A. cerana* appear similar when the community is characterized at the phylotype level, metagenomic analysis clearly demonstrates that the communities are highly distinct, being composed of divergent SDPs and strains.

### The diversity of the gut microbiota is higher in *A. mellifera* compared to *A. cerana*

As shown in **Fig. 1K**, the clustering analysis of the metagenomic ORFs resulted in a much larger number of clusters for *A. mellifera*, compared to *A. cerana*. This difference may in part be explained by *A. mellifera* housing a bacterial community composed of more SDPs. While 90% of the host-filtered reads map to the database for both hosts (**Fig. S1**), this fraction represents 12 SDPs for the five core phylotypes in *A. mellifera* (plus up to 4 non-core members), but only 7 in *A. cerana* (plus *Apibacter* sp., and occasionally *Lactobacillus kunkeei*) (**Fig. 1** and **Fig. S1**). Indeed, the number of gene clusters and the total genome assembly size per bee were both approximately twice as large for *A. mellifera* when using comparable subsets of host-filtered reads (**Fig. 2A** and **Fig. S6**). However, the sharp drop in the number of clusters from 95% to 90% sequence identity observed only for *A. mellifera* (**Fig. 1K**) indicates that strain-level diversity is also a contributing factor.

**Figure 2.**
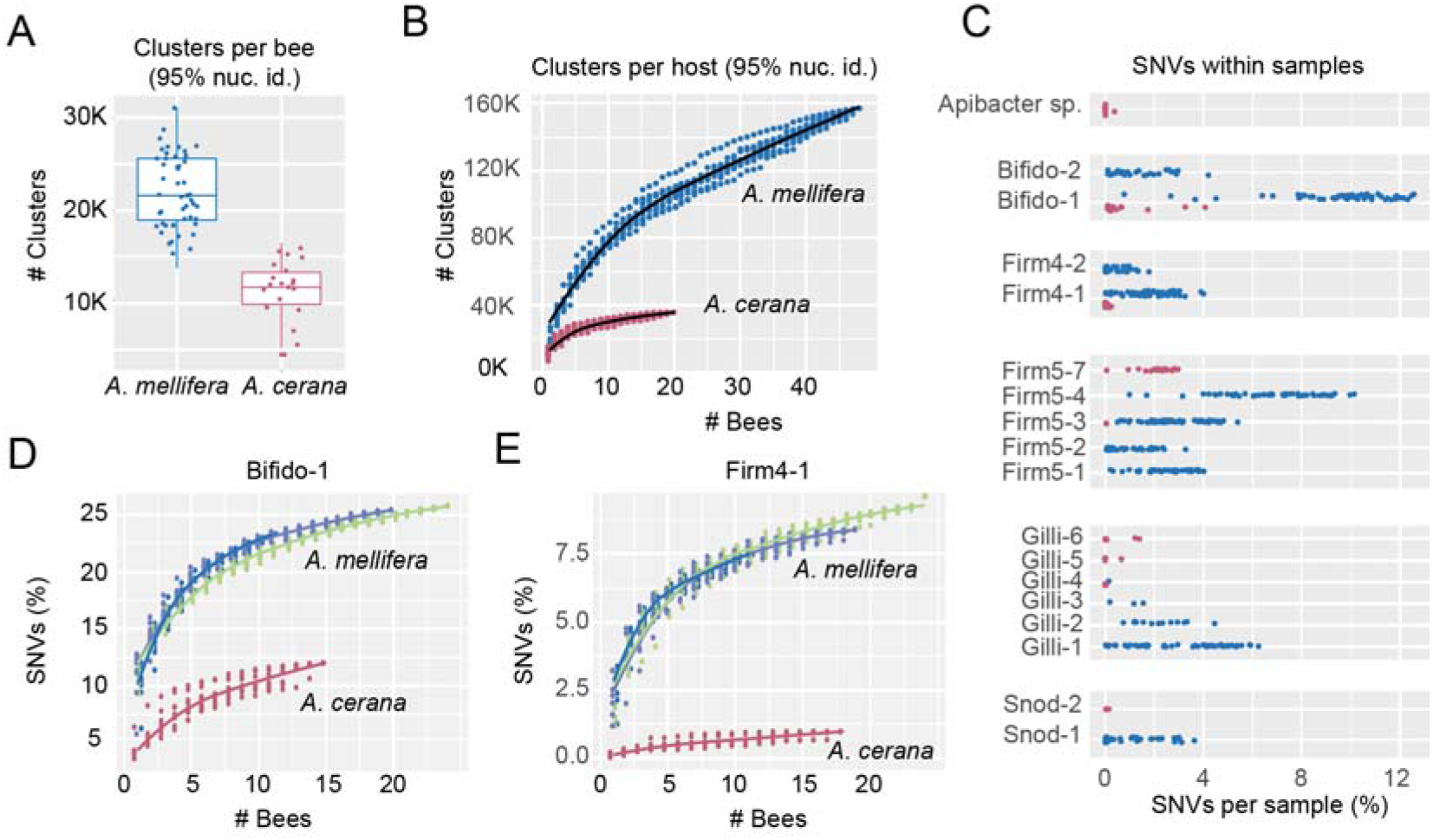
Major differences in strain-level diversity in the gut microbiota of *A. mellifera* and *A. cerana*. **(A)** Number of sequence clusters per sample for each host, based on metagenomic ORFs from assemblies generated with 20 million paired-end host-filtered reads. **(B)** Cumulative number of sequence clusters for each host, relative to the number of bee samples. **(C)** Fraction of single nucleotide variants (SNVs) within core genes in each sample, for SDPs corresponding to core phylotypes, plus *Apibacter* sp.. SDP labels correspond to Fig. 1. **(D-E)** Cumulative fractions of SNVs within core genes relative to the number of samples, for the “Bifido-1” and “Firm4-1” SDPs. Blue-green shades represent different *A. mellifera* colonies, whereas the *A. cerana* samples were pooled due to the smaller number of samples. See **Fig. S8** for curves corresponding to other SDPs and mappings to alternative reference genomes.

Previous analysis of strain-level diversity in *A. mellifera* showed that strains tend to segregate among individuals within colonies [6], in contrast to the SDPs, which mostly co-exist (**Fig.1D-F** and **Fig. S5**). Therefore, sequence clusters occurring only in a subset of bees are likely to represent strain-level diversity. To determine how the sequence clusters distribute across bees, we plotted the number of clusters relative to the number of samples, using the 95% nucleotide sequence identity threshold. From the cumulative curves (**Fig. 2B**), it is evident that the gene content harbored within individual bees represent a minor fraction of the total gene content present across hosts. Moreover, the number of sequence clusters increased more rapidly with sample size for *A. mellifera* as compared to *A. cerana*, and did not appear to have reached saturation with the current sampling size. This difference was not related to diversity between colonies or countries for *A. mellifera* (**Fig. S7**). Rather, the gene content of the gut microbiota is highly variable among bees, also within colonies, and much more so for *A. mellifera* compared to *A. cerana*. Taken together with the consistent taxonomic profile among bees (**Fig. 1D-F** and **Fig.S5**), these results indicate that a major fraction of the variation in gene content is related to strain-level diversity.

To quantify the extent of strain-level diversity, the total fraction of single nucleotide variants (SNVs) within core genes was determined for all SDPs. At the level of individual bees, only two of the SDPs colonizing *A. cerana* were found to harbor more than 2% SNVs in any of the samples (“Firm5-7”, “Bifido-1”) (**Fig. 2C**). In contrast, nearly all the SDPs colonizing *A. mellifera* had more than 2% SNVs per bee in a major fraction of the samples (**Fig. 2C**). For example, “Bifido-1”, an SDP shared between both bee species, had on average 9.6% SNVs per individual bee in *A. mellifera*, while in *A. cerana* the average percentage SNVs was as low as 0.8%.

To quantify diversity at the colony level, we again generated cumulative curves of SNVs as a function of the number of analyzed bees (**Fig. 2D-E** and **Fig. S8**). Interestingly, the two SDPs occurring in both hosts, “Bifido-1” and “Firm4-1” (**Fig. 1G and H**), displayed a clear difference in strain-level diversity between the host species (**Fig. 2D-E**). This difference was not explained by the choice of reference genome, since the pattern persisted after swapping the reference genome with an isolate from the alternate host (**Fig. S8).**

Based on these results, we conclude that strain-level diversity in *A. mellifera* is substantially higher than in *A. cerana*, both in individual bees and within colonies.

### The gut microbiota of *A. mellifera* encodes more diverse enzymes for polysaccharide degradation

The observed differences in SDP composition and strain-level diversity raise the question whether the gut microbiota is functionally distinct between the two hosts. To address this question, we annotated metagenomic ORFs using the COG and CAzyme databases. Furthermore, the analysis was restricted to the Japanese metagenomes (20 samples per host), using metagenome assemblies with 20 million host-filtered paired-end reads per sample, in order to facilitate quantitative comparisons.

Although the number of ORFs per sample was approximately twice as high for *A. mellifera* compared to *A. cerana*, the relative COG profiles were indistinguishable among hosts (**Fig. 3A**). Consistent with previous studies [26], carbohydrate metabolism and transport” (COG category “G”) was abundant across the metagenomic samples of both host species. In order to identify possible differences within this important functional category, metagenomic ORFs encoding glycoside hydrolases (GHs) and polysaccharide lyases (PLs) were annotated with dbCAN2 [27], and the number of annotations per family were counted for each sample. Notably, when calculating the mean number of annotations per family for each host species, the relative abundance of each GH/PL family was highly similar between the hosts, as evidenced by the linear correlation between the mean counts per family (**Fig. 3B**). However, samples from *A. mellifera* harbored approximately twice as many genes per family, as evidenced from the slope of the correlation (**Fig. 3B**).

**Figure 3.**
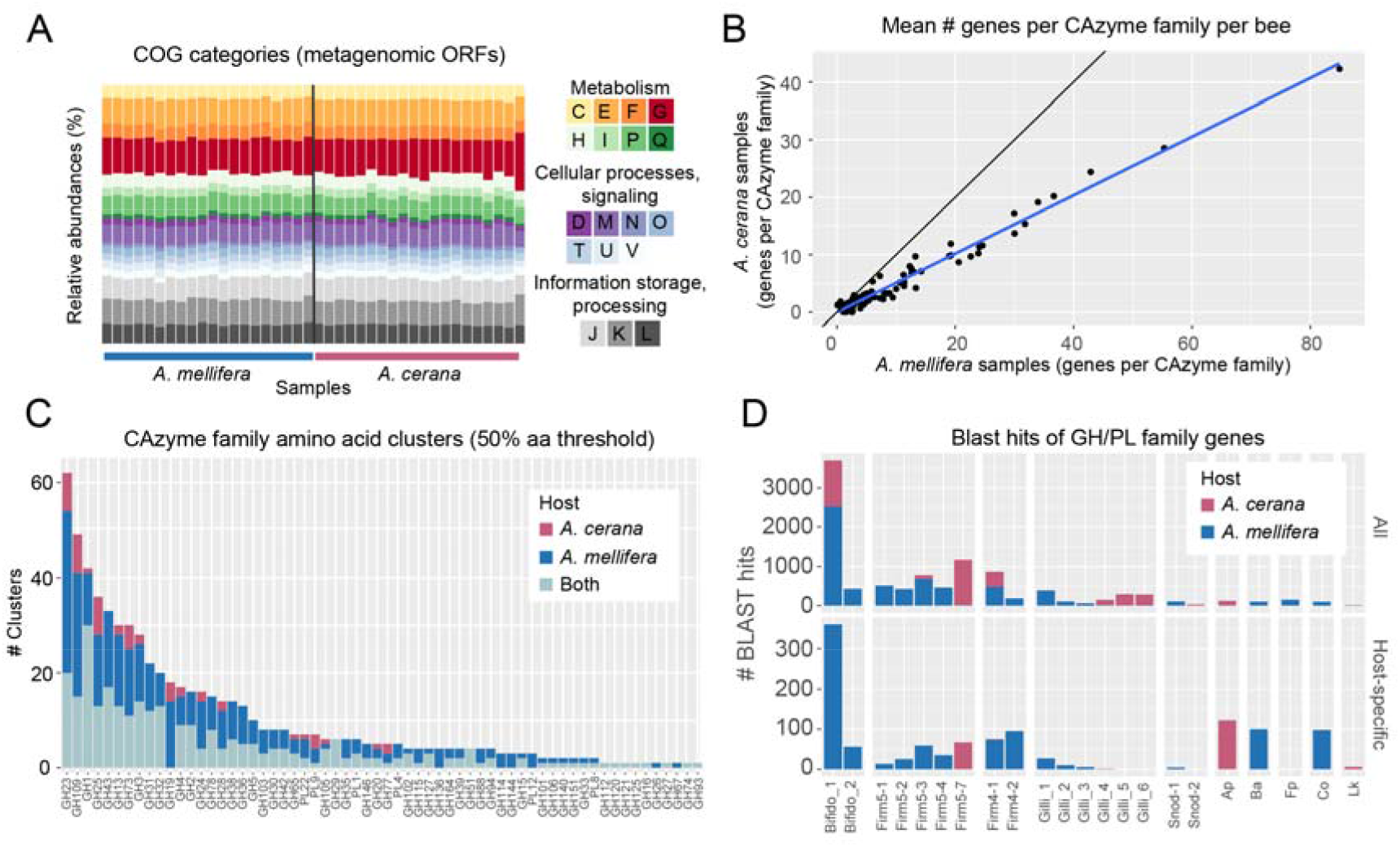
Functional comparison of the gut microbiota of *A. mellifera* and *A. cerana*. **(A)** Relative abundance of COG annotations according to general functional COG categories. **(B)** Mean number of ORFs assigned to each CAzyme family, calculated separately for each host. As shown by the blue regression line, there was a linear correlation between the counts, indicating that the CAzyme families display similar relative abundance patterns in both hosts. But, the total number of ORFs assigned to each CAzyme family is larger in *A. mellifera* samples as compared to *A. cerana* samples as indicated by the deviation from the black line. **(C)** Number of sequence clusters within each CAzyme family (clustered at 50% amino acid sequence identity), with colors indicating the subsets of clusters specific to each host, and clusters containing ORFs derived from both hosts. **(D)** SDP affiliation of blast hits for all ORF sequencess annotated as GH/PL CAzyme families, in the honey bee gut microbiota database. Results are only shown for close hits (>95% amino acid sequence identity). The upper panel includes all ORFs passing this threshold, the lower panel includes the subset of these occurring within host-specific clusters. All analyses were carried out on only the Japanese metagenome samples, to facilitate quantitative comparisons (20 samples per host, with assemblies based on 20 million paired-end host-filtered reads per sample). SDP labels correspond to Fig. 1. Species abbreviations: Ap - *Apibacter* sp., Fp - *Frischella perrara*, Co: *Commensalibacter* sp., Lk - *Lactobacillus kunkeei*.

Although the annotation of GH/PL families is based on sequence homology, a wide range of specificities have been reported within families [28]. Therefore, to estimate the diversity among the genes within GH/PL families, they were clustered separately for each family, using a highly conservative threshold (50% amino acid sequence identity). Even at this threshold, the clustering resulted in an average of 10.4 clusters per family, indicating that a major fraction of the genes annotated to the same family is likely to have distinct functions or substrate specificities. A total of 52 out of 62 GH/PL families contained host-specific clusters (**Fig. 3C**). However, only 17 families contained clusters specific to *A. cerana*, whereas all 52 contained clusters specific to *A. mellifera* (**Fig. 3C**). Moreover, the mean number of host-specific clusters per family was 6.0 for *A. mellifera*, but only 3.1 for *A. cerana*. Taken together, these results therefore indicate that *A. mellifera* has a much larger host-specific repertoire of GH/PL families as compared to *A. cerana*.

In order to gain further insights into the origin of the GH/PL families, all sequences were blasted against the honey bee gut microbiota database. Overall, 97% of the sequences had significant hits to the database (e-value < 10e-05, > 80% query coverage), with 79% having a close hit (>95% amino acid identity). Among the sequences with close hits, the vast majority of hits were to genomes of the “Bifido-1” SDP, followed by other SDPs of the *Lactobacillus* Firm4 and Firm5 phylotypes (**Fig. 3D**, upper panel). For the host-specific GH/PL clusters, only 52% of the sequences falling within *A. mellifera*-specific clusters had a close hit to the database, whereas 67% of the *A. cerana*-specific clusters had close hits. Thus, although the current database contains fewer genome isolates from *A. cerana* compared to *A. mellifera*, it is more representative of *A. cerana* in terms of GH/PL families, likely as a consequence of GH/PL genes being at least partly associated with strain-level diversity. Among the sequences corresponding to host-specific GH/PL clusters, the majority of blast hits were once again to “Bifido-1” (**Fig. 3D**, lower panel). Strikingly, they all originated from *A. mellifera* metagenomes, suggesting that GH/PL families encoded by “Bifido-1” in *A. cerana* represent a subset of those present in *A. mellifera*. A similar pattern was observed for “Firm4-1”, the other SDP shared between the two host species. While GH/PL family genes matching “Firm-4” sequences were found in the gut microbiota of both honey bee species, almost all host-specific sequences came from *A. mellifera*. Instead, for the clusters specific to *A. cerana*, most of the hits were to *Apibacter* sp. and *Lactobacillus* “Firm5-7”, suggesting that diversity occurring at the phylotype- and SDP-level contribute more to functional specialization than strain-level diversity in *A. cerana*.

In conclusion, despite the similarity in the general functional profiles, *A. mellifera* harbors a much more diverse repertoire of GH/PL families, with many more host-specific GH/PL clusters compared to *A. cerana*. Moreover, the majority of GH/PL sequences were associated with the “Bifido-1” SDP, which is much more diverse in *A. mellifera* compared to *A. cerana*, suggesting that strain-level diversity in *A. mellifera* is a major contributor to functional differences between the two host species.

### *A. mellifera* and *A. cerana* differ in bacterial community size within individual bees

According to neutral theory, diversity is expected to correlate with habitat size and population size. Therefore, we sought to determine whether *A. mellifera* and *A. cerana* differ in terms of the spatial niche they provide for their gut bacterial communities. Based on wet-weight, the hindgut, where most of the bacteria reside, was not significantly different between the hosts (**Fig. 4A**). However, the bacterial community size, as estimated from quantitative real-time PCR with universal 16S rRNA primers (normalized to the copy number of the host gene actin) was still found to be significantly larger for *A. mellifera* compared to *A. cerana* (p < 0.001, Mann-Whitney U test) (**Fig. 4B**).

**Figure 4.**
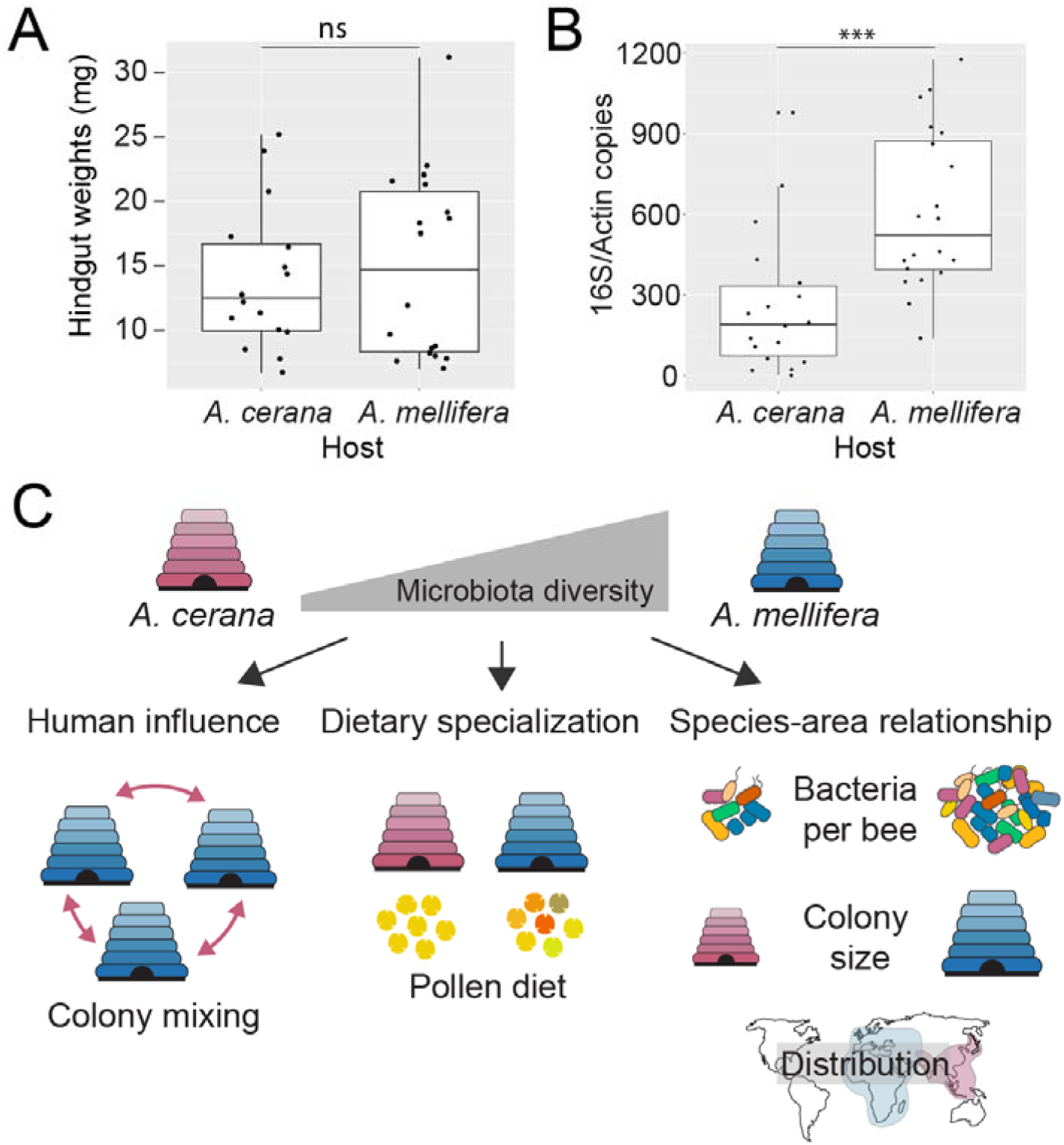
Possible factors explaining the difference in gut microbiota diversity between *A. mellifera* and *A. cerana*. **(A and B)** Wet-weight of the hindgut (where most of the bacteria reside) and qPCR results for estimation of bacterial community size (targeting the 16S rRNA gene, and normalized by copy-number of the host gene actin). Statistical significance was calculated using a Mann-Whitney *U* test (ns - not significant, ***p<0.001). **(C)** Schematic illustration of three possible factors explaining differences in diversity. “Human influence”: transportation and mixing of *A. mellifera* colonies and genotypes around the world by beekeepers, resulting in mixing of strains from different geographic origins, and thereby increasing strain-level diversity in *A. mellifera*. “Dietary Specialization”: *A. mellifera* may have a more generalist diet (here illustrated by pollen grain diversity) as compared to *A. cerana*, and therby be able to sustain a more diverse community. “Species-Area relationship”: Although previously applied to species-level diversity in animals, this concept may also apply to strain-level diversity in bacteria. For the honey bee gut microbiota, spatial differences are applicable at three levels: The size of the bacterial community within individual bees, the size of honey bee colonies, and the size of the geographic range.

## Discussion

In the current study, we carried out a community-wide metagenomic characterization of the gut microbiota of two closely related honey bee species, *A. mellifera* and *A. cerana*. From this analysis, three key results emerged. Firstly, we found that the gut bacterial communities of the two host species were highly divergent, consisting of different SDPs and strains, despite having a very similar phylotype-level composition. Secondly, the two host species displayed major differences in the magnitude of strain-level diversity within their bacterial communities. And thirdly, the gut microbiota of *A. mellifera* harbored a much larger repertoire of enzymes related to polysaccharide breakdown. Thus, in the time since their last common ancestor, approximately 6 million years ago [20, 23], the gut bacterial communities of *A. mellifera* and *A. cerana* seem to have undergone substantial changes in composition, genomic diversity, and functionality, with likely consequences for the interaction with their hosts.

Based on amplicon sequencing of the 16S rRNA gene, multiple studies have shown that gut microbiota composition is influenced by host phylogeny [29, 30] with the overall observation that closely related host species harbor more similar gut bacterial communities at the phylotype-level than more distant ones [31]. However, the slow evolutionary rate of the 16S rRNA gene [9] does not permit evolutionary analysis of phylotypes that are shared across related hosts. Therefore, there is currently little data providing insights into the evolution of the gut microbiota for closely related animal hosts. Targeting the fast-evolving *gyrA* gene for three bacterial families colonizing hominids, Moeller *et al.* found evidence of co-diversification in the Bacteroidaceae and Bifidobacteriaceae, but not the Lachnospiraceae [32]. Similarly, for honey bees and bumble bees, amplicon-sequencing of the *minD* gene uncovered both host-specific and more generalist clades within the core phylotype *Snodgrassella* [19]. Consistently, comparative genome analyses of bacterial isolates have also uncovered several examples of host-specific lineages [14–17]. However, the current study is the first to report community-wide patterns of host specialization using metagenomic data.

Interestingly, while we found evidence of host-specialization for all of the five core phylotypes colonizing both hosts, the extent of divergence differed widely among them. For three of the phylotypes (*Lactobacillus* Firm5, *Gilliamella*, and *Snodgrassella*), each host was found to be colonized by different SDPs, i.e. bacterial lineages which are sufficiently divergent to be classified as different species [18]. In contrast, the host-specialization of the *Bifidobacterium* and *Lactobacillus* Firm4 phylotypes only became evident when analyzed with strain-level resolution, consistent with the comparatively short branch lengths observed for the core genome phylogenies. Contemplating a 16S rRNA gene divergence rate of about 1% per 50 million years [33, 34], it seems unlikely that any of the SDPs could have emerged within the 6 million years separating *A. mellifera* and *A. cerana*. More likely, the current SDP composition represents a selection of pre-existing SDPs, with secondarily evolved traits resulting in the currently observed host preference. For example, we found that all SDPs within the *Lactobacillus* Firm5 phylotype contributed to highly divergent host-specific glycoside hydrolases, which could potentially allow for dietary specialization and host adaptation. In contrast, the comparatively little sequence divergence between the host-specialized strains of *Lactobacillus* Firm4 and the Bifidobacteria could match a time span of 6 million years, and thus be a product of co-diversification. However, a larger genomic dataset from multiple host species will be needed to properly test this hypothesis.

Remarkably, we also found that the two host species substantially differed in terms of the extent of genomic diversity in their gut microbiota. In comparison, a previous study employing amplicon-sequencing of the 16S rRNA gene found only a subtle trend towards lower diversity in *A. cerana* compared to *A. mellifera*, even when using a highly elevated sequence identity clustering threshold of 99.5% [13]. Given that the 20 metagenomic samples of *A. cerana* came from only two colonies in Japan, it remains to be confirmed whether our findings also apply to other subspecies of *A. cerana* outside of Japan. However, considering that (i) the phylotype composition of the samples in the current study was largely consistent with the previous 16S rRNA-based results for *A. cerana* [13], (ii) no geographic patterns were observed for *A. cerana* in the previous 16S rRNA-based study, which included a broad sampling across Asia and Australia [13], and (iii) no quantitative differences in strain-level diversity was observed across colonies and continents for *A. mellifera* in the current study, it seems unlikely that the strain-level diversity should differ strongly across colonies, at least from a quantitative perspective. Given that many studies have shown a lack of concordance between the 16S rRNA gene and genome-level divergence in bacteria [9, 11, 35], also in the honey bee gut microbiota [36], these results seem to rather highlight that genome-wide data is needed for quantification of strain-level diversity, especially at short evolutionary time-scales. This is also supported by the findings from the amplicon-sequencing of the *minD* gene, which revealed marked differences in strain-level diversity for the phylotype *Snodgrassella* between bumble bees and honey bees, suggesting that the 16S rRNA gene strongly underestimates diversity in the honey bee gut microbiota [19].

However, despite a rapid increase in metagenomic studies during the last decade, quantification of strain-level diversity from metagenomic data is still technically challenging and quantitative comparisons across hosts are therefore rare. Differences in sampling, DNA extraction, and sequencing depth can have a strong impact on both composition and diversity, making cross-comparisons between samples and studies particularly difficult [37]. For example, for the human gut microbiota, it was estimated that a cumulative genome coverage of at least 1000x would be required to obtain a representative sampling of the strain-level diversity [38], a number which is still well beyond the current norm. Even in simple communities, like the bacterial endosymbionts colonizing deep-sea mussels, quantification of strains can be sensitive to sequencing depth [4]. In the current study, technical biases were limited by using a common sampling and DNA extraction protocol, and a comparable sequencing depth. Given the simple taxonomic composition of the honey bee gut microbiota, we were able to obtain deep sequencing of the individual community members, with comparable mapping and *de novo* metagenome assembly efficiencies in both hosts. Furthermore, normalization by sampling and sequencing depth was applied in all analyses. Taken together, we are therefore confident that the observed community-wide differences in strain-level diversity are not due to technical biases.

Several factors could potentially have contributed to the observed difference in strain-level diversity (**Fig. 4C**). Firstly, although *A. cerana* has long been used for honey production in Asia, *A. mellifera* differs from all other extant honey bee species by having been extensively transported around the globe by humans, for hundreds of years [25]. Thus, it is possible that humans have contributed to a mixing of locally adapted strains, thereby increasing diversity within colonies (**Fig. 4C**, “Human influence”). As of yet, large-scale studies on the distribution of strains are still lacking for honey bees, and previous studies have reported mixed results [13, 19]. Likewise, in the current study, ordination plots based on shared SNVs clustered by country for only a subset of SDPs. Future studies on wild honey bee populations should provide further insights into this question. Interestingly, if the high strain-level diversity in *A. mellifera* is caused by human interference, it raises the possibility that the gut microbiota of *A. mellifera* is sub-optimal for local conditions, potentially resulting in reduced colony fitness.

However, based on molecular data, *A. mellifera* has had a large and varied geographic range long before human interference [20, 21]. Moreover, despite the occurrence of colony collapse disorder in managed colonies around the world [39–41], *A. mellifera* has been found to succesfully establish feral colonies even outside its native range (i.e. the New World), indicative of a remarkable ability to survive under highly varied conditions [42]. Thus, it is possible that *A. mellifera* is more of a generalist than *A. cerana*, for example by having a wider foraging range, and therefore is able to maintain a more varied bacterial community and larger repertoire of sugar breakdown functions (**Fig. 4C**, “Dietary specialization”). Indeed, diet has been shown to have an impact on diversity in the gut microbiota in multiple studies, with for example a reduction of diversity reported for the human gut microbiota as a consequence of westernized diet [43, 44]. As of yet, large-scale systematic studies on the foraging preferences of *A. cerana* and *A. mellifera* have not been conducted, but they appear to have largely overlapping foraging ranges in Asia [45, 46]. Further studies are therefore needed to determine whether differences in strain-level diversity are related to dietary differences.

Finally, it is also possible that the difference in strain-level diversity between *A. mellifera* and *A. cerana* is driven by neutral processes (**Fig. 4C**, “Species-area relationship”). Specifically, the “species-area relationship” posit a positive correlation between habitat size and diversity, and is widely held to constitute one of the few laws in ecology [47–49]. However, it is still debated whether this relationship also applies to bacteria [50–52]. The gut microbiota represents an attractive model system for testing the hypothesis, since bacterial populations in this case can be easily delineated, due to the host-association. Indeed, it has previously been proposed that differences in gut microbiota diversity in different bee species could be explained by the species-area relationship, since lower 16S rRNA gene diversity was found to be correlated with both smaller colony size and smaller bacterial population size within individual bees across honey bees, bumble bees, and stingless bees [13, 19]. Interestingly, our results uncovered much more dramatic differences in diversity at the strain-level, compared to the species level, raising the question of whether the “species-area” relationship must be adapted to include strain-level diversity in bacterial communities. In the case of *A. mellifera* and *A. cerana,* which are very similar in terms of colony cycle and behaviour, spatial differences also exist at multiple levels possibly explaining the differences observed in strain-level diversity (**Fig. 4C**, “Species-area relationship”). Firstly, we found that the size of the bacterial communities within individuals was significantly larger for *A. mellifera* compared to *A. cerana*, a trend which was also observed in the previous study comparing diversity across bees [13]. Secondly, *A. mellifera* is known to form larger colonies than *A. cerana* [22]. And thirdly, the native range of *A. mellifera* is also larger compared to *A. cerana* [25, 53, 54]. Our analysis clearly shows that the diversity within individual bees is much lower than the diversity found within their colonies. Thus, it seems plausible that competition among strains is alleviated when strains colonize different hosts. If so, it also follows that larger colonies should be able to support more strain-level diversity, by providing more colonization opportunities for strains that would otherwise compete. However, given that the diversity was also consistently higher for individual bees of *A. mellifera* as compared to *A. cerana*, the species-area relationship may be equally applicable at this level. In contrast, the inclusion of multiple colonies had very little impact on diversity, suggesting that host geographic range is of less importance for maintenance of strain-level diversity in honey bees.

In conclusion, the results of the current study brought several fundamental questions regarding the evolution and maintenance of diversity in host-associated bacterial communities to the foreground. While the term “diversity” has an inherently positive connotation, it is not obvious whether diversity in host-associated bacterial communities should be beneficial, and if so, in what sense [55]. For example, high strain-level diversity in the gut microbiota of *A. mellifera* may provide more metabolic flexibility, facilitating foraging on more diverse pollen sources, and thereby faster adaptation to changing environmental conditions. On the other hand, high strain-level diversity could also lead to increased competition within the gut microbiota, with resources being diverted towards inter-bacterial warfare, rather than host-symbiont mutualistic interactions. It is also possible that a less diverse gut microbiota consisting of strains adapted to local conditions would be more beneficial than a more diverse one. These possibilities can be experimentally tested in honey bees, thereby providing novel insights into the functional relevance of strain-level diversity in host-associated bacterial communities and for honey bee health.

## Methods

### Bioinformatic pipeline: custom scripts and intermediate result files

All bioinformatic tasks were done with custom scripts, using perl, bash or R, unless otherwise indicated. In order to facilitate the revision, we provide access to all databases and scripts used, via the following link: https://drive.switch.ch/index.php/s/W24Ay35HSSTLHkZ. The complete pipeline will be made publicly available on Zenodo upon publication of the peer-reviewed article.

### Metagenomic sampling and sequencing

A total of 40 metagenomic samples were collected from individual bees, with 20 samples from *A. cerana* and *A. mellifera*, respectively. For each host species, 10 worker bees were collected from each of two colonies, soaked in absolute ethanol and stored at −20 °C until dissection. All sampled colonies were from different apiaries (“AIST”, “UT”, “Chiba”, “Hodogaya”) located less than 100 km apart, close to Tsukuba, Japan, in September 2017. DNA was extracted from each gut as described previously [6] and sequenced using the NexteraXT library kit, with paired-end sequencing (2 x 100bp), on an Illumina HiSeq2500 instrument. The quality of the raw data was checked with fastQC [56], and the reads were subsequently trimmed with Trimmomatic [57] using the settings: LEADING:28, TRAILING:28 MINLEN:60, and trimming for the Nextera adapter. The same filtering was also applied to 36 previously published metagenomes [6], corresponding to age-controlled bees (Day 10 and Day 22/24), collected from the apiary at the University of Lausanne, Switzerland [6].

### Quantification of bacterial loads

In order to estimate the size of the bacterial communities colonizing the gut of *A. mellifera* and *A. cerana*, 8-10 honey bees were collected from each of two colonies, for each host. For *A. mellifera*, samples were collected in May 2019, from the same apiaries as the metagenomic samples (but corresponding to different colonies). For *A. cerana*, samples were collected in September 2017 from Hodogaya (same colony as for metagenomic samples) and in October 2017 from NIES (different apiary). Dissected guts were placed in bead-beating tubes (2ml) with 364 µL ultra-pure DNase/RNase-free ddH_2_O and zirconia/silica beads, and stored at −80 °C until DNA extraction. DNA was extracted using a CTAB-based protocol [58]. For each sample, 364 µL of CTAB lysis buffer (4% CTAB (w/v), 0.2M Tris-HCl pH8.0, 2.8M NaCl, and 0.04M EDTA pH8.0), 2 µL of 2-mercaptoethanol, and 20 µL of 20 mg/mL proteinase K was added. Bead-beating was done twice for 90 s, at 3.5 krpm (MicroSmash MS-100), with 1 min. rest on ice in between. 1 µL of 10mg/mL RNase A was added, and the tubes were incubated overnight at 55 °C. Next, 750µL PCI (phenol-chloroform-isoamyl) was added, the samples were mixed by shaking, placed on ice for 2 min, and centrifuged at 13.3 krpm at 4 °C for 30 min. DNA was precipitated with ethanol, washed, air-dried and dissolved in 50µL ultra-pure DNAase/RNase-free ddH_2_O.

Bacterial loads were estimated with quantitative real-time PCR, targeting the V3-V4 region of 16S rRNA gene with the following primers: 5’-ACTCCTACGGGAGGCAGCAGT-3’ (forward) and 5’-ATTACCGCGGCTGCTGGC-3’(reverse) [59]. Normalization was done relative to the actin gene of the host, using the following primers for *A. mellifera*: 5’-TGCCAACACTGTCCTTTCTG-3’ (forward) and 5’-AGAATTGACCCACCAATCCA-3’ (reverse). For *A. cerana,* the reverse primer was 5’-AGAATTGATCCACCAATCCA-3’. Standards were prepared as in [60]. qPCR reactions were performed in triplicates in a total volume of 10 µL, containing 5 µL of 2x TB Green premix Ex Taq II, 0.2 µL ROX reference dye II, 0.2 µM of each primer and 1 µL of 100x-diluted extracted DNA, on a QuantStudio 3 instrument (Applied Biosystems). The thermal cycling conditions were as follows: denaturation at 95 °C for 30 s, followed by 40 amplification cycles at 95 °C for 5 s, and 60 °C for 1 min (actin) or 30 s (16S rRNA).

The wet-weight of the hindguts for each host was estimated by collecting additional honey bees from AIST (*A. mellifera*, n=18, sampled in August 2019), and NIES (*A. cerana*, n=16, sampled in September 2019). The guts were dissected and weighed on the day of collection, without freezing.

### Establishment of a non-redundant genomic database

A previously published honey bee gut microbiota genomic database [6] was updated to include recently published genomes isolated from *A. mellifera*, plus genomes isolated from other social bee species. Additionally, ten new genome isolates from *A. cerana* were sequenced with PacBio to increase the database representation for this host species for the current study. Pairwise average nucleotide identities (ANI) were calculated with fastANI [18] for all isolates of the same phylotype, and used to streamline the database for redundancy. Genomes isolated from *A. mellifera* or *A. cerana* were required to have a maximum of 98.5% ANI to other genomes within the database, whereas genomes isolated from other species were streamlined at circa 95% ANI, in both cases prioritizing the most complete genome assemblies. Metadata for all genomes included in the database is provided in Dataset S1.

### Metagenomic assemblies

To filter off host-derived reads, the reads of each metagenomic sample were first mapped against a database containing the genomes of *A. cerana* (PRJNA235974) and *A. mellifera* (PRJNA471592, version Amel_Hav3.1), using bwa mem [61] with default settings. Bam files containing unmapped reads were generated with samtools [62] (flag -f 4), and paired reads were extracted from the bam files with Picard tools [63]. Additionally, subsets of the host-filtered reads were generated in increments of 10 million read pairs for each sample. Metagenomic assemblies were generated independently for each sample and each read subset, using SPAdes [64] with default settings for metagenomic assembly. The resulting contigs were filtered to have a minimum length of 500bp and a minimum kmer coverage of 1 (parsed from the contig fasta header). To check for eventual differences in assembly efficiency related to community complexity, the reads of each subset were mapped back to the corresponding assembly with bwa mem, and the number of mapped reads was calculated with samtools (flag -F 4).

### SDP metagenomic validation

An overview of the pipeline used for SDP validation is given in Supplementary Figure S8. Candidate SDPs were identified using core genome phylogenies (Supplementary Figure S3a: schematic illustration of 4 candidate SDPs, from two phylogenies) and pairwise average nucleotide identities (ANI), as described previously [6]. Validation was done separately for each candidate SDP. In step 1, alignments of the core gene sequences in the gut microbiota database were generated, for all the genomes associated with the candidate SDP (Supplementary Figure S3b, illustrated by colored arrows and lines in the alignments). Furthermore, the core gene sequences were used as queries in a blastn search against a database containing all ORFs (predicted with Prodigal [65]) on all metagenomic assemblies. Sequences of metagenomic ORFs were extracted from the blast file for hits where the ORF length was at least 50% of the query length and the blast alignment identity was above 70%. In step 2, the sequence of each metagenomic ORF was added individually to the corresponding core gene alignment (using mafft [66], with the option --addfragment) (Supplementary Figure S3b, illustrated by grey arrows and lines in the alignments), and the maximum percentage identity within the alignment was recorded (using Bioperl)[67]. Additionally, the recruited metagenomic ORFs were blasted against the gut microbiota genomic database, and their closest SDP was recorded (based on blast hit percentage identity). For plotting, recruited ORFs with best hit to the SDP being evaluated were assigned to the first density distribution (Supplementary Figure S3c-d, shown in color), while recruited ORFs with a best hit to other SDPs were assigned to the second density distribution (Supplementary Figure S3c-d, shown in grey). An SDP was considered confirmed if the two distributions were non-overlapping (Supplementary Figure S3c), indicating that the metagenomic ORFs recruited to the SDP were discrete relative to related candidate SDPs contained within the database.

### Community profiling

For each sample, all quality-filtered paired-end reads were mapped against the gut microbiota genomic database, using bwa mem with default settings, and subsequently filtered using a minimum alignment length of 50bp. The unmapped reads (extracted with samtools flag -f4 and picard tools) were mapped to the host genomic database, to calculate the fraction of host-derived reads per sample.

The relative abundance of community members within samples was quantified based on filtered single-copy core genes, as described previously [6], with the only modification being that the coverage of core genes was extracted with ‘samtools bedcov’ (instead of ‘samtools view region’). Specifically, using the bam files corresponding to mapped reads only, the coverage depth per core gene region was extracted with ‘samtools bedcov’, and divided by the gene length (resulting in average mapped read depth per bp). As previously [6], the mean gene coverage was summed up per core gene family, and the abundance of the phylotype/SDP was estimated by terminus coverage, using a fitted segmented regression line.

### SNV profiling

SNVs occurring within core genes were profiled using a reduced genomic database, containing one representative genome per SDP. For the two SDPs represented by isolates derived from both *A. cerana* and *A. mellifera* (“Bifido-1” and “Firm4-1”), the pipeline was repeated using a database containing a reference genome isolated from the alternative host species, in order to check the impact of reference genome choice on SNV quantification. The reads were mapped against the reduced database (using bwa mem), and the bam-files were filtered by both alignment length (min 50bp) and edit distance (max 5). Candidate SNVs were predicted with freebayes [68] and gnu parallel [69], using the following options: “-C 5 --pooled-continuous --min-alternative-fraction 0.1 --min-coverage 10 --no-indels --no-mnps --no-complex”. Thus, complex variants were not predicted, and rare SNVs (less than 10% relative coverage within samples) were excluded. Filtering was carried out using a combination of custom perl scripts and vcflib [70]. For each SDP, only samples with at least 20x terminus coverage were profiled. Furthermore, core gene families were excluded, if they had less than 10x mean coverage in any of the profiled samples. Finally, candidate SNVs were excluded if freebayes reported “missing data” for more than 10% of the profiled samples. Only SNVs remaining polymorphic across samples after all filtering steps were included in the downstream analysis.

From the resulting filtered SNV files, the fraction of SNVs was calculated within and across all samples, cumulative curves were generated using 10 different sampling orders, and the pairwise fractions of shared SNVs were calculated (i.e. jaccard distance), as described previously [6].

### Gene content diversity and functional characterization of communities

To compare diversity and functional profiles in a quantitative manner across samples and hosts, ORFs were predicted with prodigal (option --meta) on metagenome assemblies based on 20 million host-filtered paired-end reads per sample. Predicted ORFs were further filtered to minimize the impact of spurious annotations, by excluding ORFs shorter than 300bp or flagged as partial by Prodigal (partial flag 11, 01 or 10). Sample affiliations were added to the fasta headers of the filtered ORFs, after which the sequences were concatenated.

To quantify gene content diversity within and across samples and hosts, the filtered ORFs were clustered with CD-HIT [71], using a range of thresholds (80-95% nucleotide identity), with hierarchical clustering as recommended in the manual. For each threshold, the number of shared and host-specific clusters were counted from the cluster file. To estimate the extent of segregation of sequence clusters among individuals, cumulative curves depicting the number of clusters relative to the number of bees were also calculated and plotted.

Amino acid sequences of filtered ORFs were annotated with eggnogmapper (version 1.0.3) [72, 73], from which COG category annotations were extracted and counted. Polysaccharide lyases and glycoside hydrolases were annotated using the dbCan2 database, v8 [27]. The database was queried with hmmsearch, and the results were filtered by e-value (max e-value 1e-05 for alignments longer than 80 amino acids, otherwise 1e-03), and HMM coverage (min fraction 0.3). To investigate whether the two host species contain a similar repertoire of carbohydrate-active enzymes, the mean number of annotations per CAzyme family was calculated for each host. Furthermore, in order to estimate sequence divergence and host specificity within CAzyme family annotations, ORFs were clustered separately for each CAzyme family, using CD-HIT with an amino acid identity threshold of 50% and a word-size of 3, as recommended in the CD-HIT manual. Finally, ORF sequences annotated as either glycoside hydrolases (family “GH”) or Polysaccharide lyases (family “PL”) were blasted against the honey bee gut microbiota database, in order to determine which SDPs contribute most to these enzyme families (threshold of significance: e-value < 10e-05 and query-coverage > 80%, with “close hits” corresponding to the subset of these having an amino acid percentage identity > 95%).

### Data Availability

The metagenomic data has been deposited under Bioproject PRJNA59809.

## Acknowledgments

We would like to thank Yoshiko Sakamoto, Shohei Kanari, Nao Tsuyuki, Yoshihiro Saito, Hiroki Kohno, and Takeo Kubo for their help during the sampling of honey bees. We also thank Kohei Fukuda for the technical assistance of molecular work. P.E. is supported by the European Research Council ERC-StG ‘MicroBeeOme’ (714804) and the Swiss National Science Foundation SNFS project grant 31003A_179487. R.M. and P.E. are supported by the Human Frontier Science Program HFSP Young Investigator grant RGY0077/2016.

## Contributions

P.E. K.E. and R.M. designed the study. S.S. performed the experimental work, including sample collection, DNA extraction, and qPCR. K.E. developed the bioinformatic pipeline and analyzed the data. P.E. contributed to the data analysis. K.E. and P.E. wrote the first draft of the manuscript. R.M contributed to the editing of the manuscript.

