## Supplementary Information for "Vast differences in strain-level diversity in the gut microbiota of two closely related honey bee species"

**Content:**

Supplementary Figures S1-S8
Supplementary Datasets S1-S3 (only legends, data provided as separate files)
Supplementary Material and Methods
Supplementary References

### Supplementary Figure

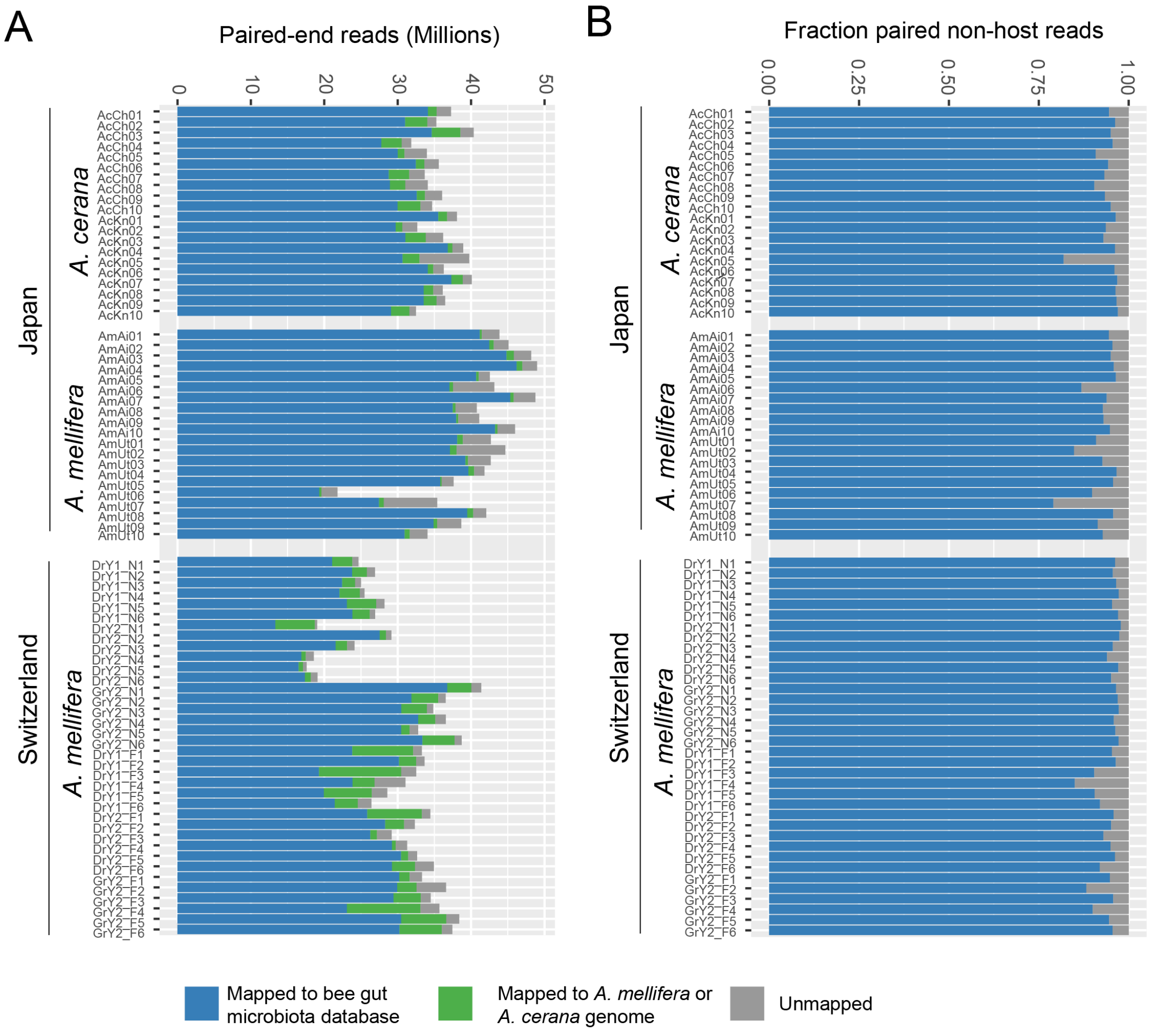


**Figure S1. Mapping results on novel genomic database. (A)** Total number of reads (unpaired) mapped to the honey bee gut microbiota database (blue), to the host genome database (green) and unmapped (grey). **(B)** The relative fraction of reads mapped to the honey bee gut microbiota database (blue) and unmapped reads (grey), excluding host-derived reads.

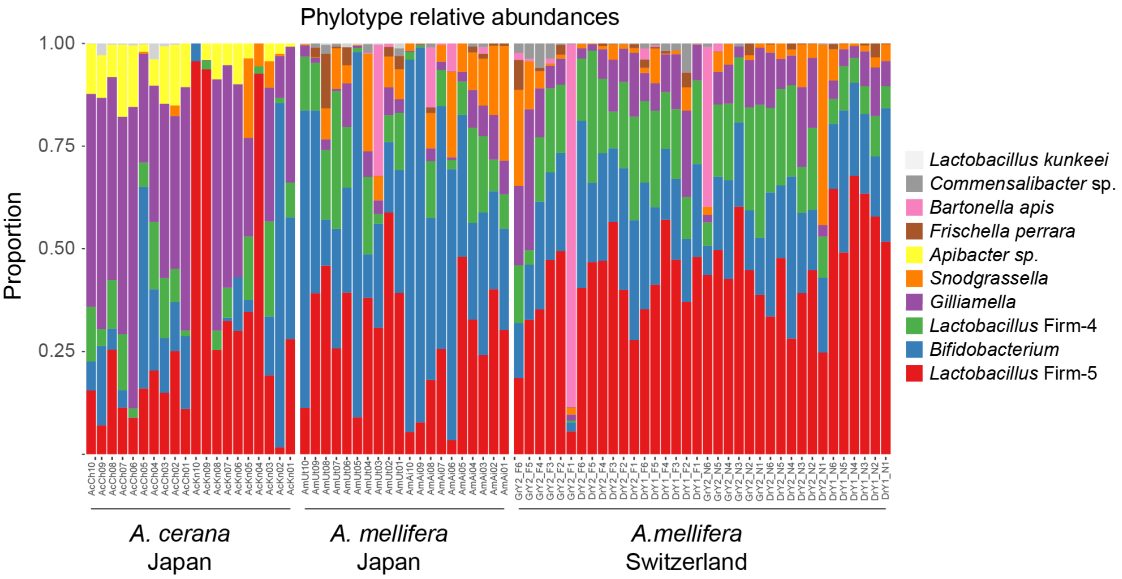


**Figure S2. Phylotype-level composition of the gut microbiota in *A. mellifera* and *A. cerana*.** The relative abundance of phylotypes was quantified based on mapped read coverage to single-copy core gene families, with a segmented regression line to estimate the genome copy-number at the terminus of replication. Sample names are the same as in **Fig. S1**. The host and country affiliation of the samples are shown below the graphs.

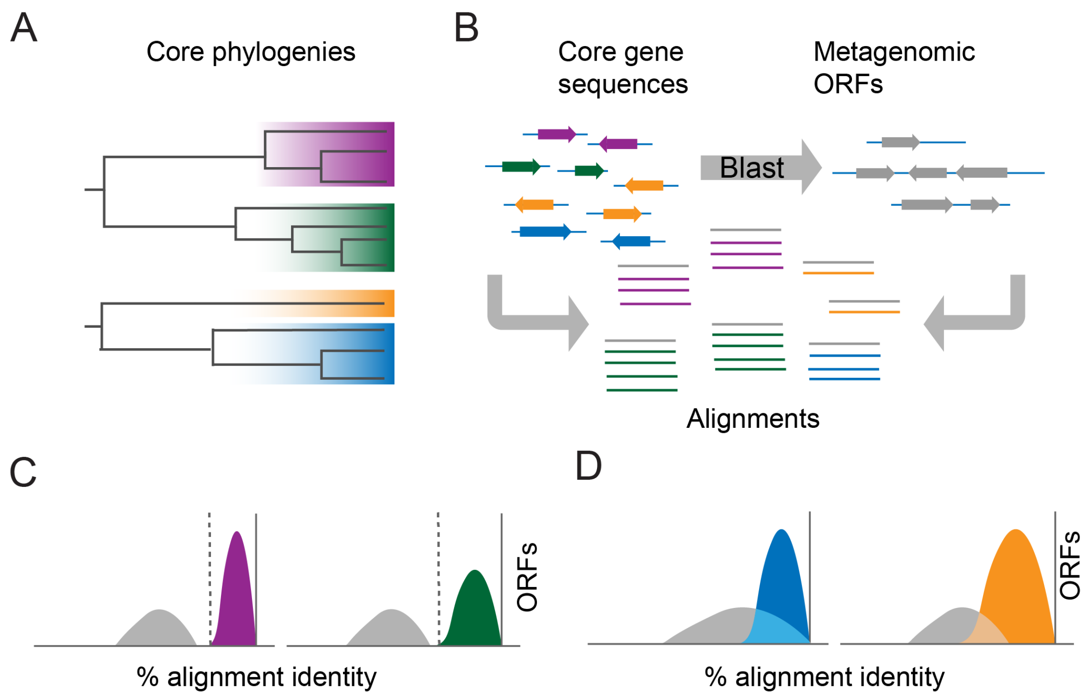


**Figure S3. Bioinformatic pipeline for inferring SDPs. (A)** Candidate SDPs are inferred from isolate genomes, based on core genome phylogenies and pairwise ANI. In this example, two schematic core genome phylogenies are shown, each with two candidate SDPs, as indicated by the color shades. **(B)** Core genes are extracted from isolate genomes (colored arrows), and aligned separately for each core gene family and SDP (colored lines). Additionally, metagenomic ORFs are recruited to each SDP by blasting the core gene sequences against the metagenomic ORFs (grey arrows). Recruited ORFs are added individually to the core gene alignments (grey lines), and their maximum percentage identity to the core genes is calculated. **(C,D)** Density distribution plots of the maximum percentage identity of all recruited ORFs. The colored distributions correspond to recruited ORFs with a best blast hit to the candidate SDP being evaluated, grey distributions correspond to recruited ORFs with a closer hit to another SDP in the database. An SDP is considered confirmed if the two distributions are largely non-overlapping **(C)**.

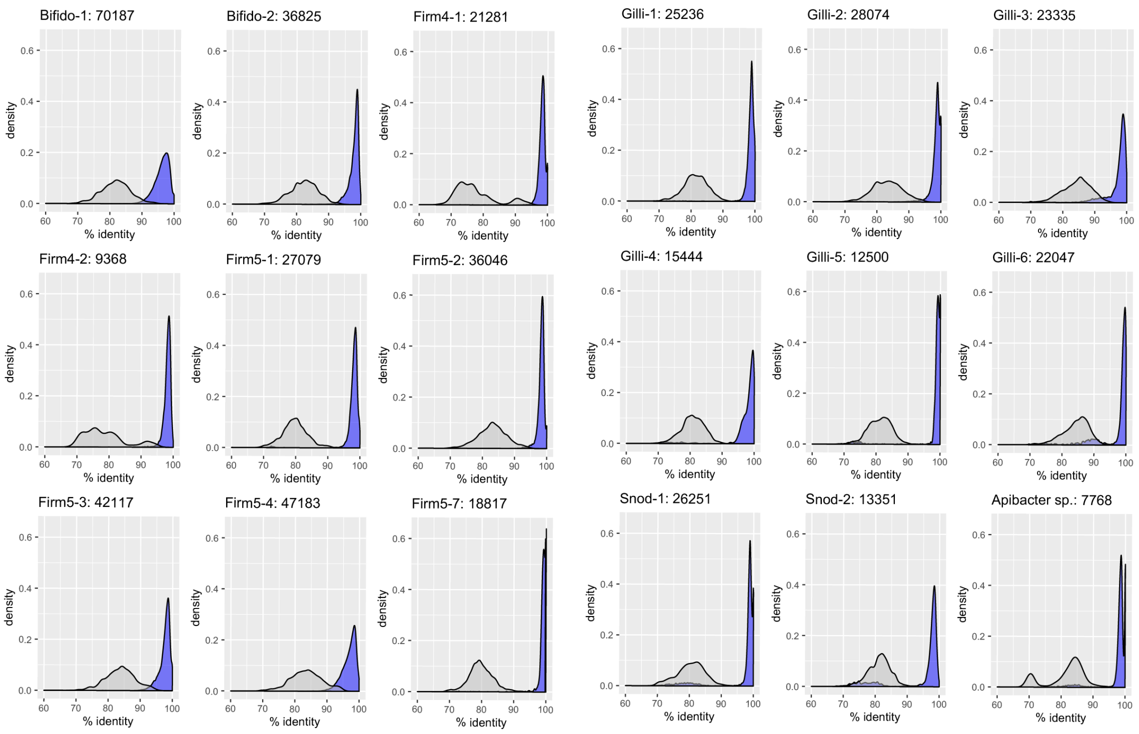


**Figure S4. Results of metagenomic validation of candidate SDPs.** The SDPs were validated by recruiting metagenomic ORFs to core gene families in the database. The number of non-redundant recruited ORFs is indicated in the panel titles of each SDP. For each recruited ORF, the maximum percentage alignment identity to the core genes within the SDP was calculated, and the distribution of all hits was plotted. Within each panel, the blue distribution represents the subset of recruited ORFs with a best blast hit to the candidate SDP, while the grey distribution represents recruited ORFs with a higher-scoring blast hit to another SDP in the database. An SDP is considered confirmed if the two distribution are largely non-overlapping (i.e. discrete relative to each other).

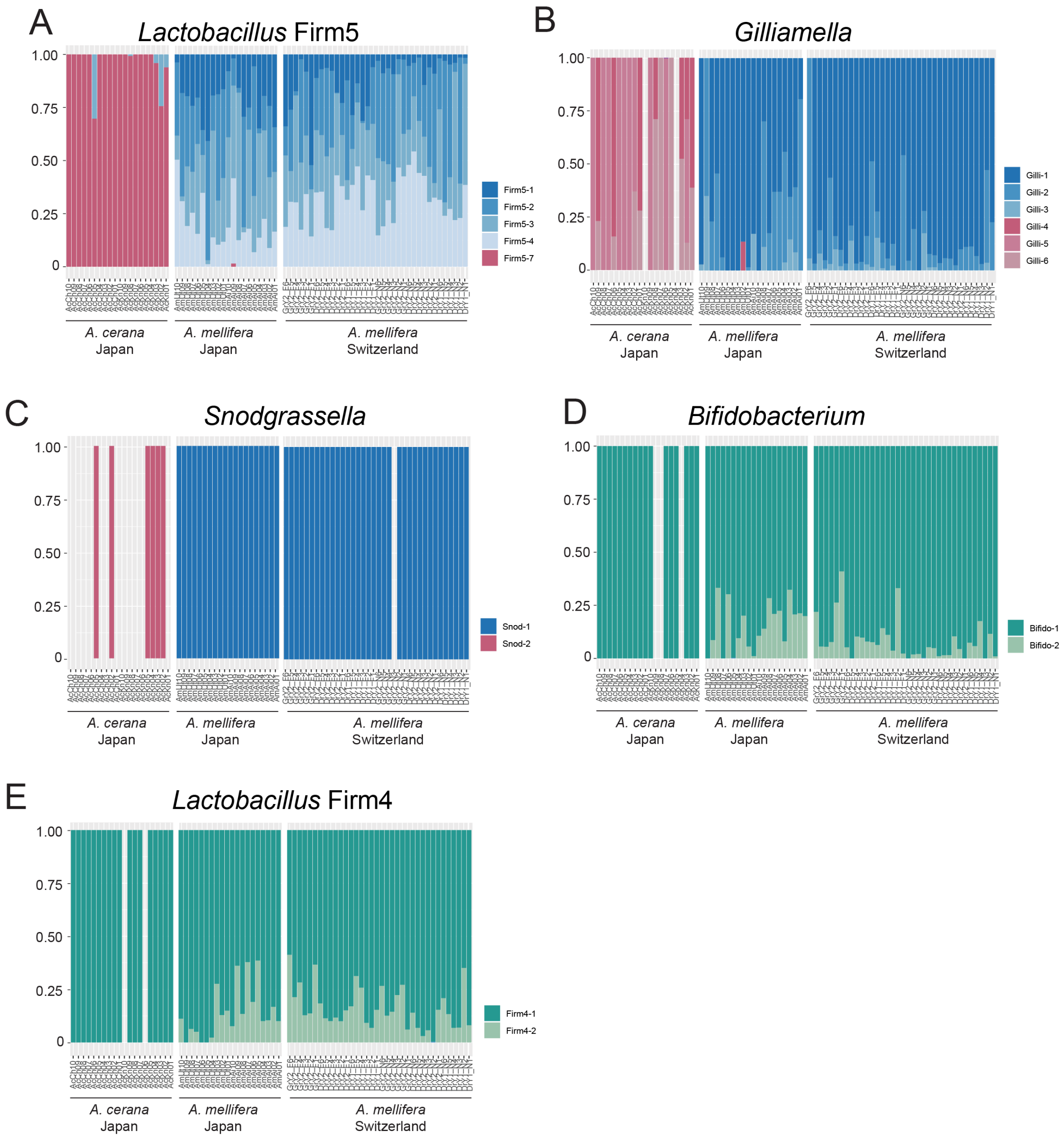


**Figure S5**. **Relative abundance of SDPs for all core phylotypes, including swiss samples.** Barplots displaying relative abundance of confirmed SDPs within all five core phylotypes colonizing both *A. mellifera* and *A. cerana*.

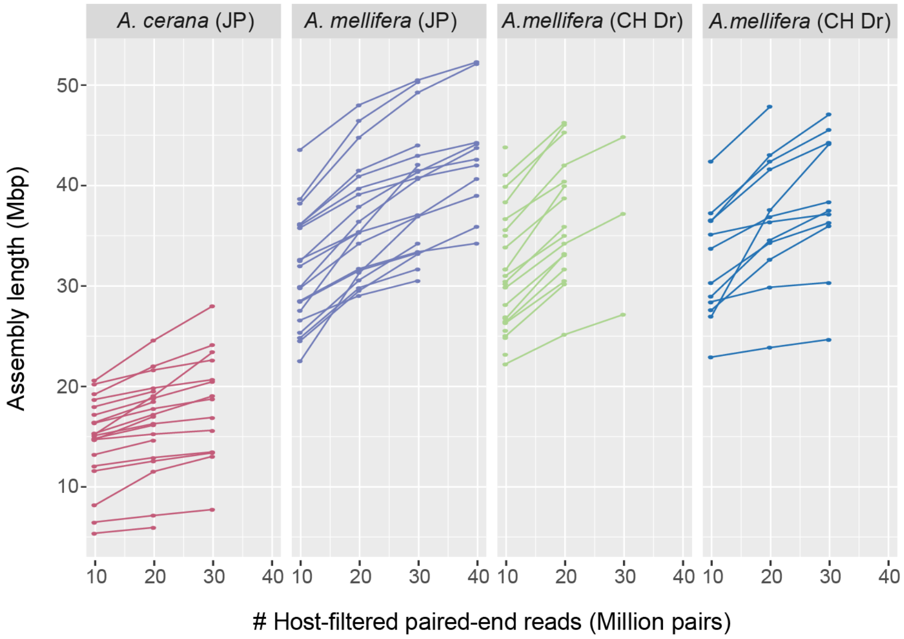


**Figure S6**. **Metagenome assembly size versus number of host-filtered reads, for different hosts and colonies.** Subsets of host-filtered reads were generated in increments of 10 million paired-end reads, and used for *de novo* metagenome assembly with SPAdes. The assembly sizes were calculated on filtered contigs (min length 500bp, min kmer coverage 1). Individual lines represent read subsets from the same sample.

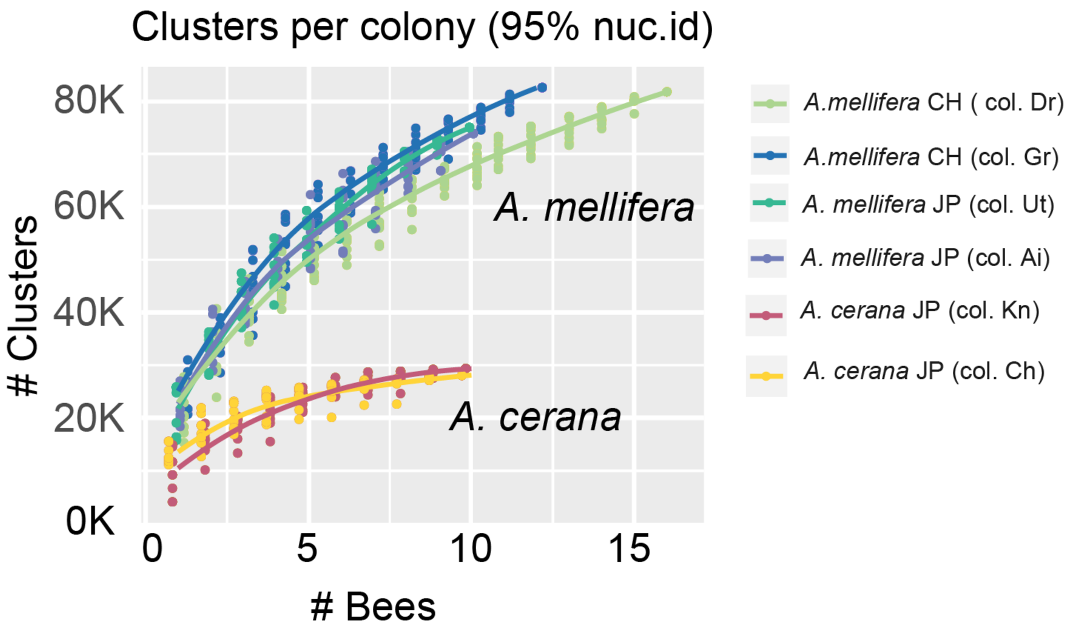


**Figure S7. Cumulative curves of the number of sequence clusters relative to the number of samples, split per colony**. Sequence clusters were generated from all metagenomic ORFs, using a nucleotide clustering threshold of 95% sequence identity. The number of clusters per colony was counted from the cluster file, skipping clusters for which the colony was not represented. Ten random sampling orders were generated per colony (with individual data points represented by dots). Blue/green colors represent *A. mellifera* colonies, while yellow/red colors represent the two *A. cerana* colonies.

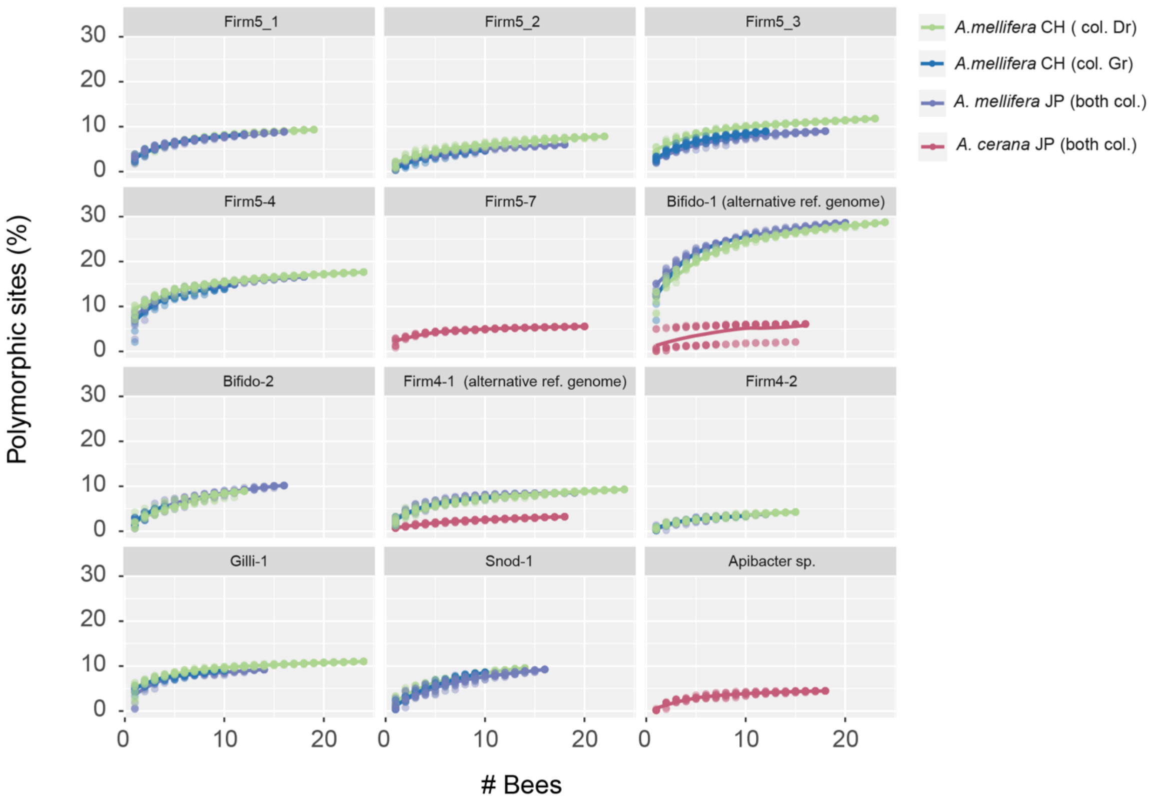


**Figure S8: Cumulative curves of the fraction of SNVs relative to the number of samples.** Curves were generated for SDPs where at least 10 samples had sufficient coverage for SNV profiling (min 20x terminus coverage), in at least one colony. Ten random sampling orders were generated per SDP (with individual data points represented by dots). Different colors represent different colonies. For “Bifido-1” and “Firm4-1”, the curves represent the SNV profiling results using a database with a genome isolate representative from the alternate host (Compared to **Fig. 2D-E**).

**Supplementary Datasets**

**Dataset S1.** Metadata for all genomes included in the honey bee gut microbiota database. For each genome, the accession numbers for both NCBI and IMG are provided (if available). Annotation files were based on the IMG files, when possible. Genomes and genes are represented by their locus-tags in the database, and throughout all analysis files. SDP names are given only when metagenomic validation has been performed.

**Dataset S2.** Pairwise average nucleotide identity (ANI) among genomes included in the honey bee gut microbiota database. Genome identifiers correspond to locus-tags, see **Dataset S1** for associated metadata.

**Dataset S3**. Principal coordinate analysis based on shared pairwise fractions of single nucleotide variants (SNVs). The distance matrix was based on the pairwise fractions of shared SNVs (jaccard distance), among all pairs of samples with sufficient coverage of SNV profiling (minimum 20x terminus coverage). Blue dots represent *A. cerana* samples, green dots represent *A. mellifera* samples from Japan, and red dots represent *A. mellifera* samples from Switzerland, with different shades indicating colony affiliation.

*SDP metagenomic validation*

An overview of the pipeline used for SDP validation is given in Figure S3. Candidate SDPs were identified using core genome phylogenies and pairwise average nucleotide identities (ANI), as described previously (1). Validation was done separately for each candidate SDP. Alignments of the core gene sequences in the gut microbiota database were generated, for all the genomes associated with the candidate SDP (Figure S3B, illustrated by colored arrows and lines). Furthermore, the core gene sequences were used as queries in a blastn search against a database containing all ORFs (predicted with Prodigal (12)) on all metagenomic assemblies (Figure S3B, grey arrows). Sequences of metagenomic ORFs were recruited to the core gene alignments when the ORF length was at least 50% of the query length and the blast alignment identity was above 70%. Recruited metagenomic ORF were added individually to the core gene alignment using mafft (13), with the option --addfragment (Supplementary Figure S3B, illustrated by grey lines in the alignments), and the maximum percentage identity within the alignment was recorded, using Bioperl(14). Additionally, the recruited metagenomic ORFs were blasted against the gut microbiota genomic database, and their closest SDP was recorded (based on blast hit percentage identity). For plotting, recruited ORFs with best hit to the SDP being evaluated were assigned to the first density distribution (Figure S3C-D, shown in color), while recruited ORFs with a best hit to other SDPs were assigned to the second density distribution (Figure S3C-D, shown in grey). An SDP was considered confirmed if the two distributions were non-overlapping (Figure S3C), indicating that the metagenomic ORFs recruited to the SDP were discrete relative to related candidate SDPs contained within the database.
