## Supplementary figures and images for "Vast differences in strain-level diversity in the gut microbiota of two closely related honey bee species"

### Supplementary Dataset S3

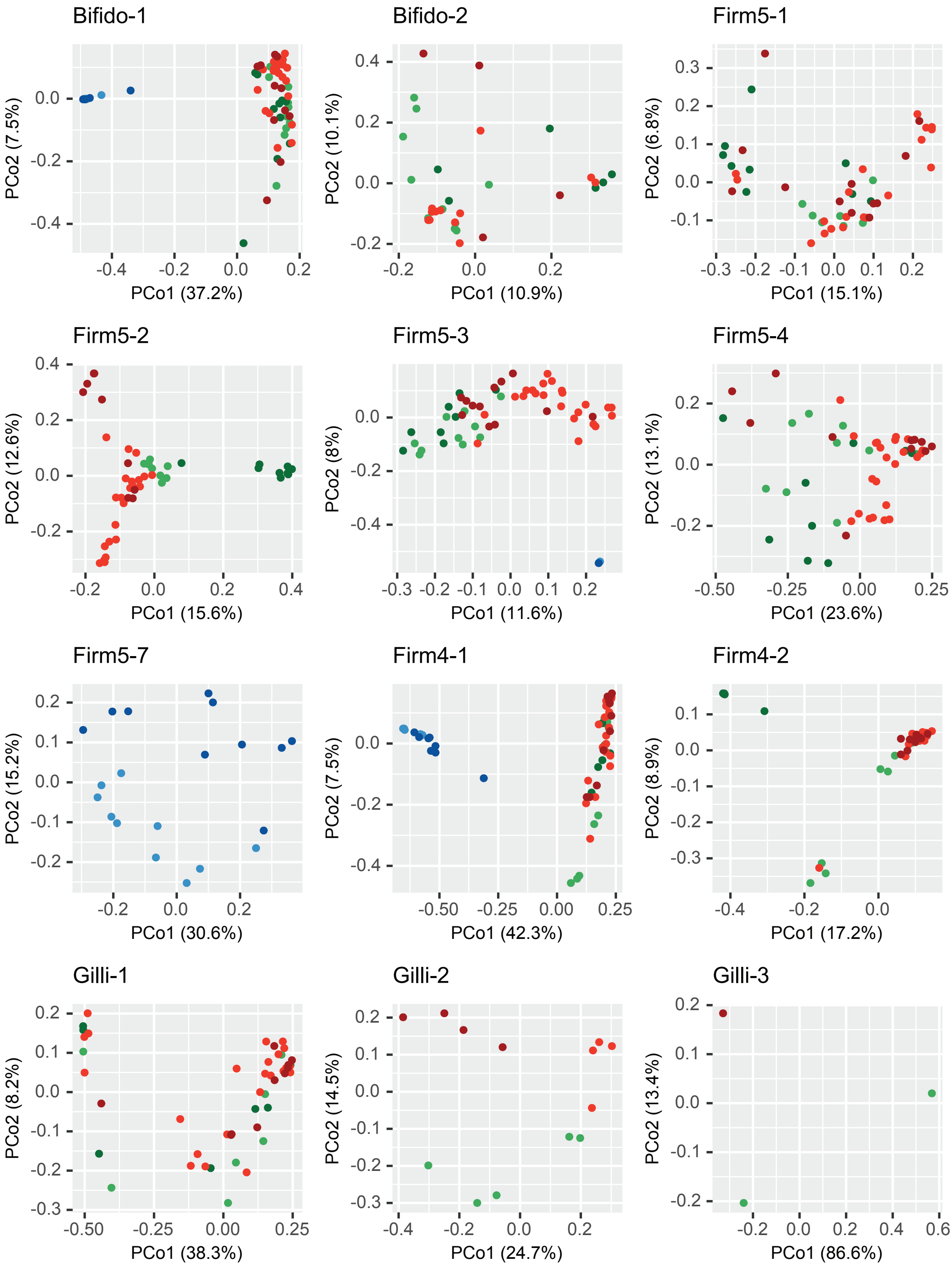

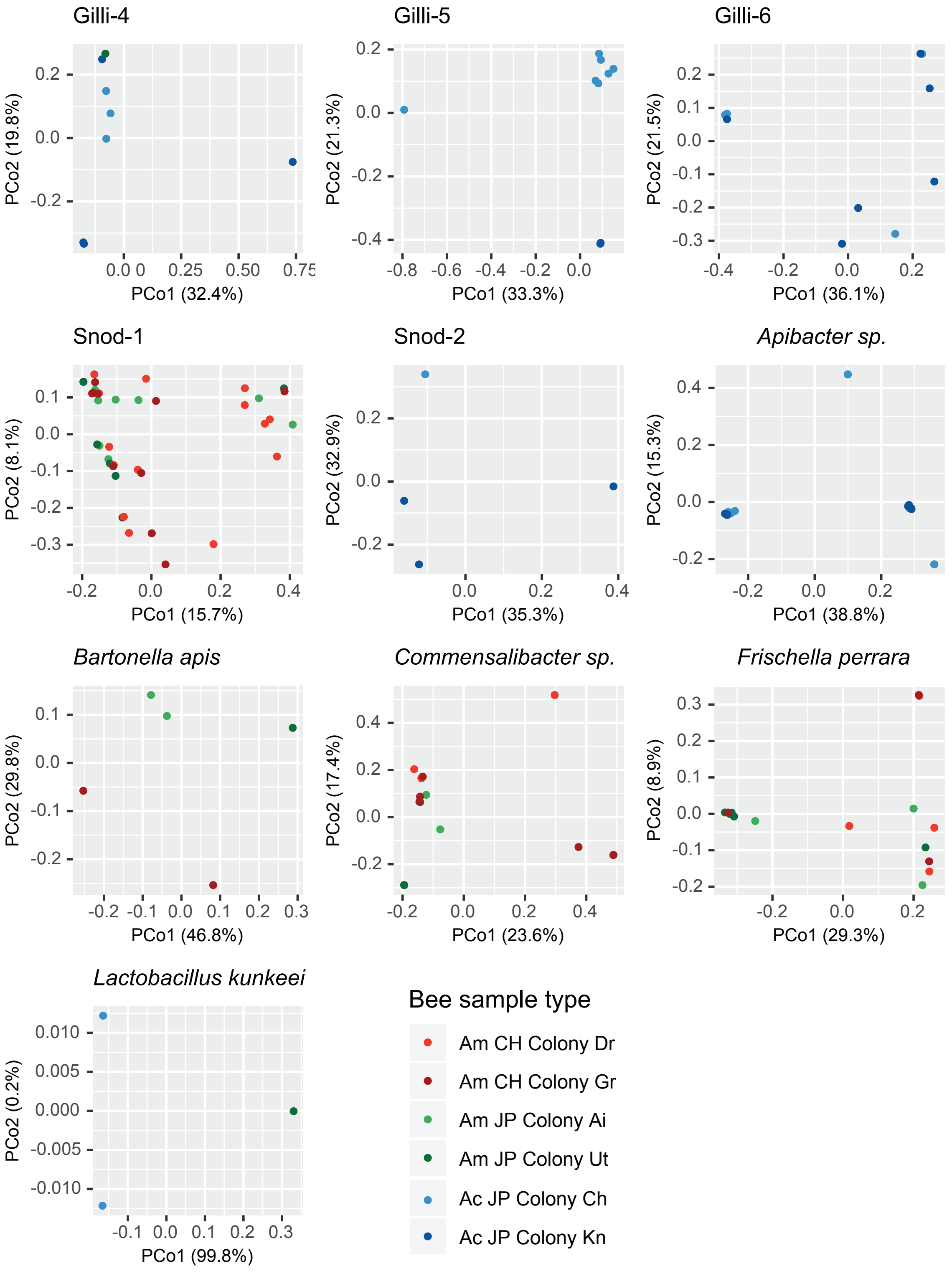
